# Ensemble kinetic modelling links residual enzyme activity to clinical symptoms in mitochondrial β-oxidation defects

**DOI:** 10.64898/2026.05.05.722902

**Authors:** Christoff Odendaal, Olga Krebs, Barbara M. Bakker

## Abstract

The mitochondrial fatty acid β-oxidation (mFAO) is an important source of energy when carbohydrate stores are depleted. It is also involved in many diseases, including inherited fatty-acid oxidation deficiencies (mFAODs). Patients with the same genetic variant often present with clinically heterogeneous phenotypes, but the mechanisms contributing to this heterogeneity are poorly understood.

To investigate the underlying pathophysiology of different mFAODs, we constructed a computational model of mFAO in human liver, based on experimentally determined enzyme kinetics. A recognised, but seldom addressed challenge in metabolic modelling is the substantial uncertainty about kinetic parameter values. Whereas experimental values of some mFAO parameters are quite reproducible, others vary by up to four orders of magnitude between different reports. To address this, we generated an ensemble of kinetic models, each with the same reaction stoichiometry and rate equations, but different kinetic parameters, sampled from distributions of literature-derived values. We also comprehensively report these values and the arguments based on which they were evaluated. The resulting models were validated against available flux data, yielding a final ensemble of 51 valid models. These models recapitulate recent findings about the accumulation of medium-chain acyl-CoAs and the concomitant depletion of free CoA (CoASH) in medium-chain acyl-CoA dehydrogenase deficiency.

We applied the ensemble to a set of known mFAODs, separating them into long-chain (LC-) and short-/medium-chain (S/MC-)mFAODs. The residual activity at which clinical symptoms are known to occur corresponded well with the residual activity in the model at which pathway flux was significantly decreased in LC-mFAODs. Residual activity in S/MC-mFAODs correlated less strongly with pathway flux, but these deficiencies did show a combined flux- and CoASH-reduction effect. This comparison is of importance to researchers and clinicians, as it identifies possible ways in which insights about one mFAOD may be applied to another based on shared biochemical properties.

**Author Summary:** When building computer models of metabolic pathways, it is typical to take the “best” experimental data and use that as input into the model. However, especially when working with human cells, ethical and practical constraints often mean that even the “best” experimental data is still subject to substantial uncertainty. We explicitly modelled the uncertainty about the inner workings of fat burning (*fatty acid oxidation*). The resulting model is known as an “ensemble”.

The ensemble predicts ranges instead of single outcomes, allowing us to assess the confidence level of our predictions. We assess a set of inherited diseases – enzyme deficiencies – simulating them at different levels of severity with the ensemble. We find that the model does a good job of predicting the severity of the deficiencies at which symptoms will occur. It also allows us to identify a key difference between two subgroups within this group of deficiencies: long-chain and medium-/short-chain, depending on the size of the fats being metabolised. The long-chain variant is predicted to correlate most straightforwardly with the severity of the deficiencies, due to its effect on energy generation. Medium-/short-chain deficiencies, in contrast, have more complex consequences.

## Introduction

The primary pathway through which fatty acids are broken down is mitochondrial fatty acid β-oxidation (mFAO). Patients with inherited, monogenetic mitochondrial fatty acid oxidation defects (mFAODs) highlight the importance of the mFAO for human physiology. More than 20 mFAODs have been identified with the main pathophysiological feature being hypoketotic hypoglycaemia and energy deficiency. Depending on the specific defect and patient-specific factors, mFAOD patients may also suffer from cardiac and skeletal muscular problems [1,2].

mFAODs, like other inborn errors of metabolism, often show poor genotype-phenotype correlation, e.g. a full loss-of-function does not necessarily predict the most severe symptoms, just as a mild loss of activity does not necessarily predict mild pathology [3]. This is because pathway perturbations can be moderated or exacerbated by other changes in the network, yielding a behaviour that is dependent not only on the original perturbation but also on the broader context [4]. For instance, individuals with medium-chain acyl-CoA dehydrogenase deficiency (MCADD) with the same gene variant on *ACADM*, the MCAD gene, who grow up in the same family can present differently [5].

We have previously built a kinetic computational model of the mFAO in human liver [6,7]. This builds on previous versions based the mFAO in rat liver [8], mouse liver [9], and mouse muscle [10]. Using these models, we have investigated inborn errors of metabolism like MCADD and multiple acyl-CoA dehydrogenase deficiency (MADD) ([8,9]), as well as age-related insulin resistance [10]. Moreover, we have used them to address more fundamental questions about the impact of enzyme promiscuity [8,11] and bistability [12]. Each time accurate kinetic parameters were sought, either from experimental literature or by parameter estimation. However, no full set of convincingly plausible kinetic parameters was available for any of the models mentioned above. Parameter uncertainty is very frequently encountered in kinetic modelling [13] and the most plausible value cannot always be identified among the options [14,15]. Moreover, while parameter uncertainty and scarcity pose a challenge in general, these issues are particularly prevalent for enzyme kinetics from humans and even more so for internal tissues, like the liver [16,17].

A common solution to the problem of parameter uncertainty is to simply choose the perceived most plausible value and discard the rest without explicit justification of the choice or reporting of the alternatives. Tsikginopoulou *et al.* [18] argue that this approach neglects a large deal of biological information: parameter uncertainty is important information. Ensemble models offer a way of explicitly accounting for the kinetic uncertainties encountered in the literature and documenting the available experimentally identified parameters comprehensively. Many variations of this approach have been proposed [15,19–25]. Typically, probability distributions of possible parameter values are derived from experimental information. Different model versions are then generated by sampling parameters from these distributions [26–30]. Some studies also include topological uncertainty by inserting the parameter combinations into various plausible network topologies [18,31–33]. The resulting ensemble of models then predicts a range of metabolic fluxes and concentrations, thereby allowing the reliability of the results to be assessed.

The importance of parameter uncertainty in the human liver mFAO model was highlighted by the observation that control shifts dramatically in the rat liver mFAO model as parameter values change: at low substrate concentration (a model parameter), the enzymes of the carnitine shuttle exert most control over the flux, while enzymes downstream in the mFAO cycle take over control at high substrate concentration [11]. Stability analysis of the rat model also revealed that the existence of two stable steady states depends on changes in the underlying parameters and boundary metabolite concentrations [12].

Confronted with a large parameter uncertainty and inspired by the approach of Tsikginopoulou *et al.* [18], we took the fixed-parameter kinetic model of the human mFAO as a starting point and varied the parameter values based on available alternatives in the literature, without varying the rate equations or stoichiometry. This is known as “ensemble kinetic modelling” [34]. We compiled a compendium of experimentally determined kinetic information and a set of mini-reviews of the underlying literature on which our parameter evaluation and weighting were based. These were used to generate 51 parameter sets reflective of the uncertainty in the parameter measurements.

This ensemble of models recapitulates our previous prediction that MCADD can cause an accumulation in medium-chain acyl-CoAs which deplete the free CoA (CoASH) pool. By exploring the differences between long-chain (LC-) and short-/medium-chain (S/MC-)mFAODs with the model, we identify LC-mFAODs more straightforwardly cause a flux impairment, while S/MC-mFAODs result in a moderate flux impairment combined with CoASH depletion. The implications for understanding mFAODs and transferring risk-stratification and treatment insights between mFAODs, are discussed.

## Results

### Workflow

Figure 2A summarizes the modelling approach that we took in this study.

Previously, we used the “most plausible” parameters to make a fixed-parameter model [6,7], whereas in the present study we combined them in an ensemble kinetic model. The differences in the approaches are summarised in Fig. 2A. First, all published kinetic parameters were retrieved from the literature and compiled in a mini-review for each enzyme (DOI: 10.15490/fairdomhub.1.assay.2137.2 on FAIRDOM Hub). Subsequently, the retrieved parameters were weighted based on the origin of the enzyme (e.g. prioritizing liver over other organs and human over other organisms), the relevance of assay conditions for the in vivo situation (e.g. prioritizing measurements that were obtained at 37 °C and pH close to 7), and completeness and clarity of the methods description. The ensemble models were based on the same model topology as the previous fixed-parameter model (Fig. 1), but in contrast to earlier work we did not choose one most plausible value for each parameter. Rather, the weighted parameters were used to construct probability distributions from which specific values were sampled for each model. 50 of these models, each based on a specific sample of parameter values, together with the default model formed the ensemble of 51 models (Fig. 2A) as detailed below.

**Figure 1.**
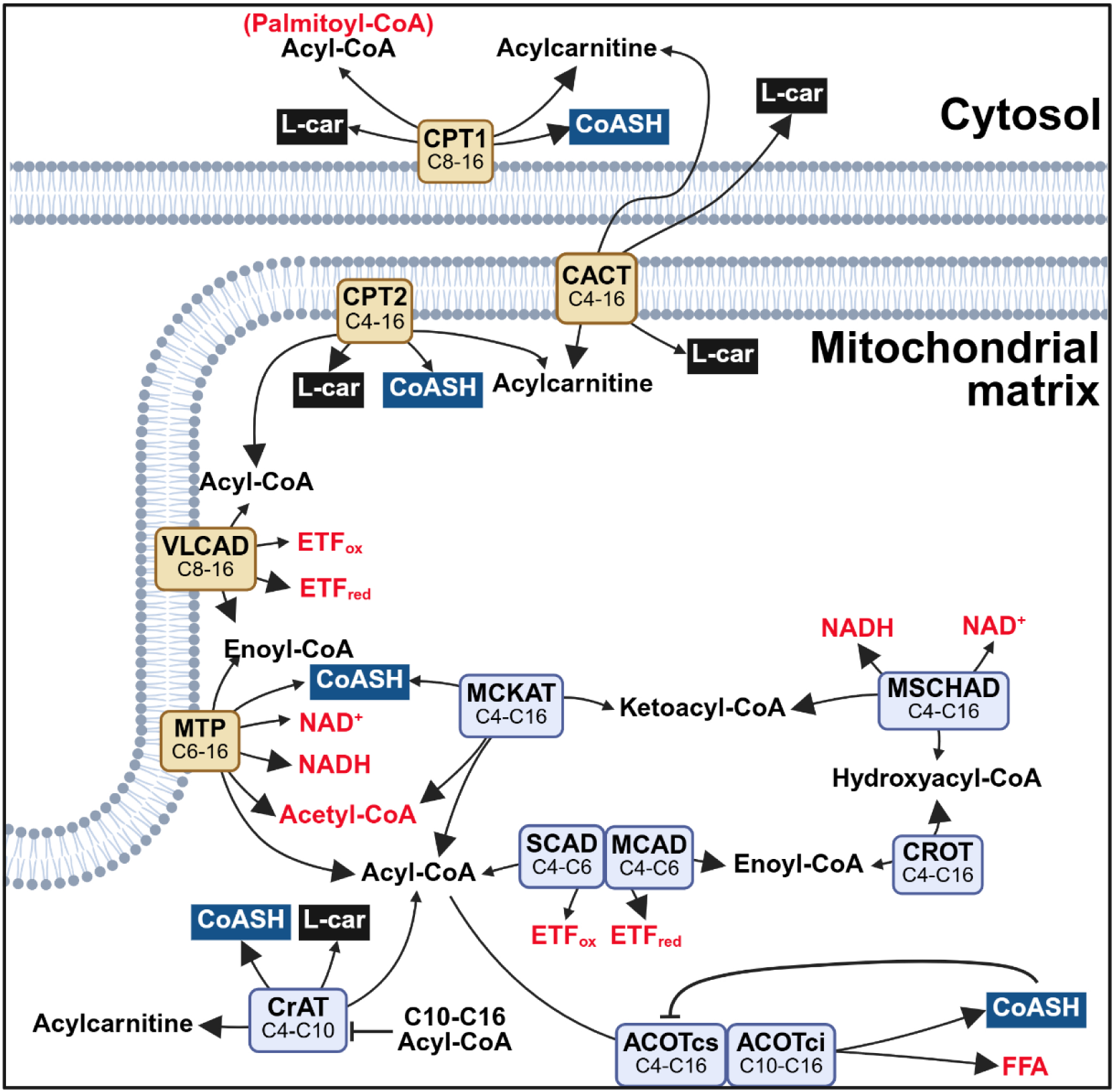
Model scheme. Metabolites in red are modelled at fixed (boundery) concentrations, the sums of L-carnitine and CoA species form two conserved moieties, respectively, on either side of the double membrane (black and dark blue boxes indicate the free moieties, L-car for L-carnitine and CoASH for CoA, respectively). All other metabolites are independent free variables. Enzymes in brown boxes are membrane-bound, enzymes in light blue boxes are matrix-localised. The longer a metabolite’s acyl chain (and hence the more hydrophobic it is) in the mitochondrial matrix compartment, the higher its concentration in the vicinity of the membrane-bound enzymes. The shorter the acyl-chain (and hence the more water soluble), the more evenly it is distributed across all enzymes in the mitochondrial matrix. All reactions are reversible, except the ACOT-catalysed reactions. Larger arrow heads indicate the forward direction in mFAO. The model structure and attributes are the same as previously published [6,7]. CPT1/2 = carnitine palmitoyltransferase 1/2, CACT = carnitine acylcarnitine translocase, CrAT = carnitine acetyltransferase, VLCAD/MCAD/SCAD = very long-/medium-/short-chain acyl-CoA dehydrogenase, MTP = mitochondrial trifunctional protein, MSCHAD = medium-/short-chain hydroxyacyl-CoA dehydrogenase, CROT = crotonase, MCKAT = medium-chain ketoacyl-CoA thiolase, ACOTci/cs = CoA-sensitive/CoA-insensitive acyl-CoA thioesterase, FFA = free fatty acid.

The variation in parameter values retrieved from the literature was substantial, often surpassing an order of magnitude (Fig. 2B and 2C). In exceptional cases as much as 10 000-fold differences were found. The size of the variation differed between parameters, with some yielding more consistent measurements than others (Fig. 2B). The weighted values and their literature sources are presented in a summarised format in Text S1.

**Figure 2.**
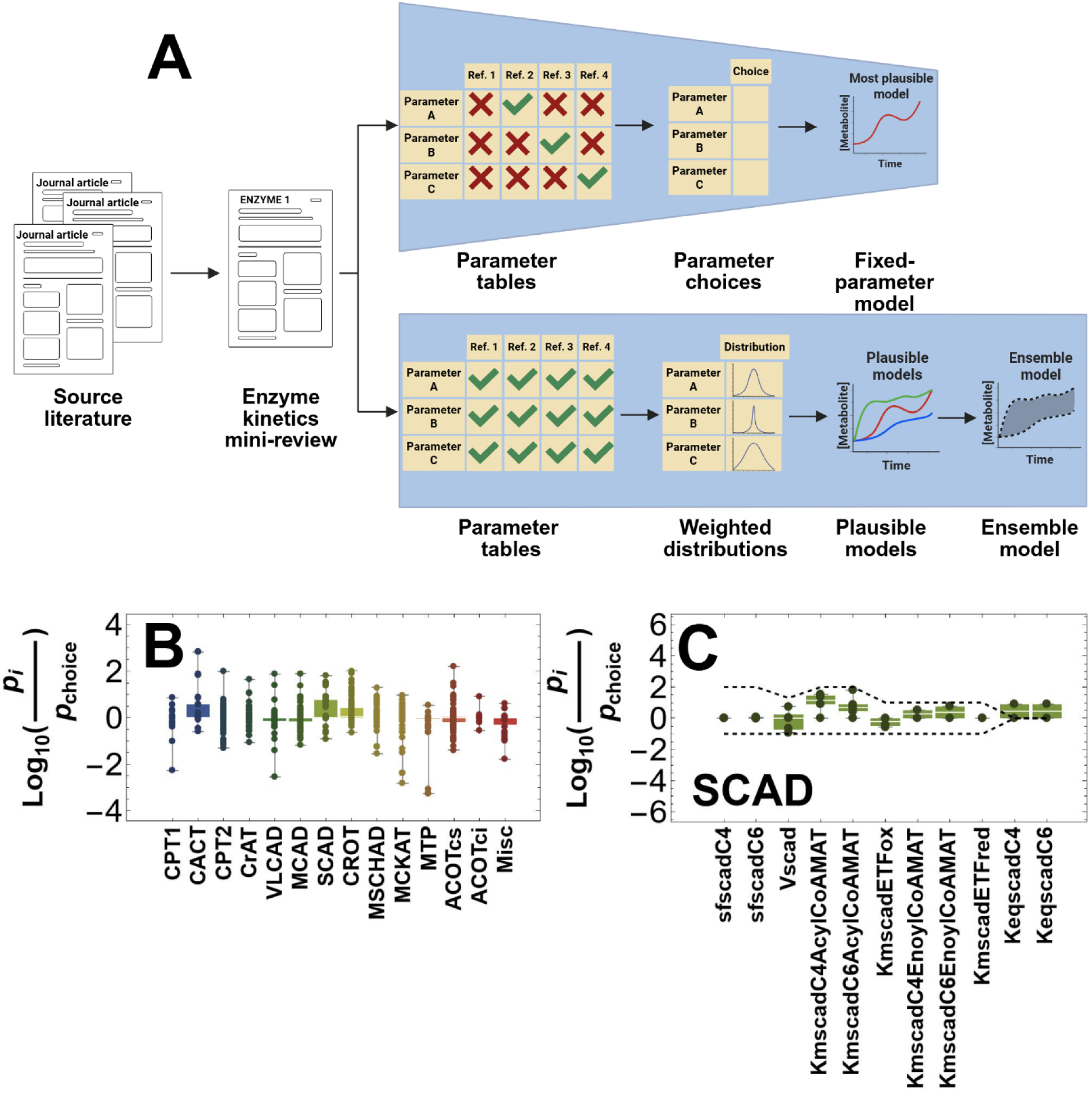
Modelling parameter uncertainty. **A.** Comparison of the fixed-parameter approach to an ensemble kinetic approach. After reviewing the literature, we previously selected the most plausible value for each parameter to make a fixed-parameter model (*top*, [6,7]). In the present study, all retrieved parameters were weighted to construct distributions and generate multiple plausible models, which together form an ensemble (*bottom*). **B.** Log_10_ of the fold difference between the alternative parameter values (*p_i_*) and the most plausible value in the fixed parameter model (*p_choice_*), including all parameters of a specific enzyme. **C**. Log_10_ of the fold difference between the alternative parameter values and *p_choice_*, for individual parameters of the enzyme SCAD. The dashed lines represent the upper and lower bounds imposed on the parameters during sampling. The per-parameter variation is displayed for each enzyme in Fig. S1-S14.

### Parameter sampling

Most parameters were assigned a probability distribution. Fig. 3 shows representative examples of different sampling approaches. The first and most straightforward approach was used for parameters that were not chain length-dependent, such as many V_max_ and K_i_ values. These were directly passed to distributions. Where enough credible values could be retrieved from the literature, log-normal distributions were constructed from which parameter values were sampled (Fig. 3A). Parameters were weighted according to *a priori* biochemical reasoning. Where only one credible value could be found, a normal distribution was used to mimic a likely measurement error (Fig. 3B). Uniform distributions were constructed where no likeliest value was forthcoming, so that a range of possible values could be randomly sampled (Fig. 3C).

**Figure 3.**
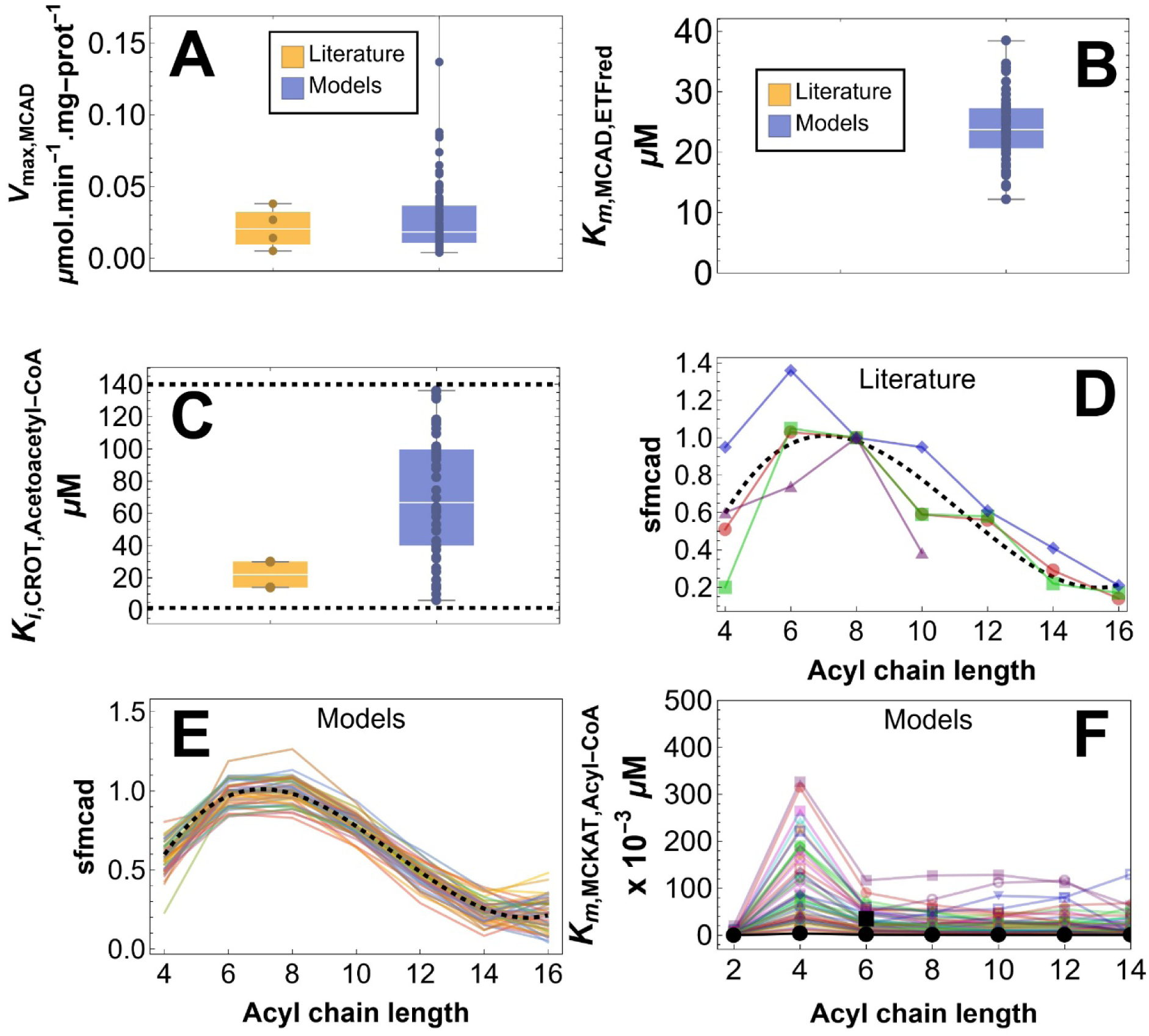
Parameter sampling examples. Typical examples of the different sampling approaches applied in this study. **A.** Parameters sampled from a log-normal distribution (blue). Four different literature values for V_max,MCAD_ (yellow) were used to construct the distribution. **B.** Parameters sampled from a normal distribution (blue). One literature value for the K_m_ of MCAD for reduced ETF was found (yellow). This value was taken as the mean and a standard deviation of 25% of the mean was assumed. **C**. Uniform distribution. Literature parameters for K_i,CROT,Acetoacetyl-CoA_ (yellow) were unsatisfactory, so a broad range of values would be investigated. A uniform distribution (between the dashed lines) was appropriate for this and yielded the sample shown in blue. **D & E.** Multivariate distribution for chain length-specific parameters. **D** shows the four literature-derived sets of specificity factors for MCAD (red, green, blue, purple) and the cubic polynomial fit to them (dashed line) with acyl chain-length as independent variable. **E** shows the same polynomial (dashed line) and 50 parameter sets (coloured lines) generated using the covariate matrix of the fitted function. **F.** Calculated parameters. The K_m_ of MCKAT for its acyl-CoA product is calculated per chain length using the sampled parameters and the Haldane relation (see text). The figure displays the calculated values as a function of acyl chain-length.

Acyl-chain length-dependent values were considered together. For instance, the specific activity of MCAD varies as a function of acyl chain length, with the highest activity observed towards medium chains (hexa- and octanoyl-CoA). This was observed in four independent studies which reported different values but showed similar patterns (Fig. 3D). The same holds for the other enzymes’ specificity factors (sf) and K_m_ values. These are a function of the length of the acyl chains, and we kept this dependence intact when fitting functions to these data (Fig. 3D). These functions and their covariance matrices were used to construct a multinormal distribution from which chain length-dependent sets could be generated (Fig. 3E). This constrained parameter variation to the observed patterns of chain length-specificity.

Three parameters (MTP and MCKAT’s K_m_ values with respect to their acyl-CoA substrates and the reverse V_max_ CACT) were calculated using the Haldane relation [35–37]. These sampled values showed very large variation (Fig. 3F), as they compound the variation of all other parameters used to calculate them. Yet, in practice this constrained the enzymes’ degrees of freedom via their equilibrium constants (K_eq_). E.g. for MCKAT:

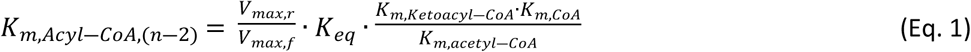

Since normal distributions technically stretch from −∞ to ∞, and log-normal distributions from zero to ∞, top and bottom bounds were also imposed on the parameter ranges to prevent unreasonable values from making it through the sampling phase. These bounds were usually 10-fold below and above the value selected for the fixed-parameter model [6,7]. The bounds imposed during sampling are explicitly indicated in Text S1 and visually represented in Fig. S1-14.

Boundary metabolite concentrations, compartment volumes, and the parameters of ACOT form the interface of the core mFAO model with its surrounding networks and were kept constant in all iterations of the ensemble. Thus, only parameter uncertainty within the mFAO is explored. Equilibrium constants are predicted with increasing accuracy based on known molecular properties of the reaction (cf. [38,39]) and were not varied in the ensemble.

### Model selection

An ensemble of 585 models with different parameter values was generated by random sampling of the parameter sets. Two experiments were simulated with each of these models, both of which were previously also used to validate the fixed-parameter model [7]. New models were generated and tested until 50 models were accumulated that satisfied all selection criteria. Including the default model, containing the literature-derived parameters, gave 51 validated models. Those models that predicted outcomes within the range of measured values for all experiments, were accepted.

The first validation experiment measured a whole-body ketogenic flux by administering a combination of stable isotope-labelled substrates to fasted adult humans [40]. Ketogenesis takes place in the liver and has acetyl-CoA as its first substrate [41]. Since the approximate proportion of mFAO-derived acetyl-CoA that enters ketogenesis is known [42], as well as the proportion of ketogenic acetyl-CoA derived from mFAO [43], the liver mFAO fluxes predicted by the ensemble of models, can be converted to ketogenic fluxes. Conversions were performed as previously [7].

The experiment was simulated at various cytosolic palmitoyl-CoA concentrations (Fig. 4A). As it is impossible to tell what the palmitoyl-CoA concentration in hepatocytic cytosol was in the whole-body ketogenesis experiment, it was decided that if a model at predicted a ketogenic flux within the range of measurements at any of the simulated palmitoyl-CoA concentrations, it was accepted. Out of 585 models generated, 274 satisfied this criterion. All rejected models predicted fluxes below the minimum of the measured range; none were rejected for passing above the maximum. Some models reached the range at intermediate palmitoyl-CoA concentrations but sank back down beneath it at higher substrate concentrations. These were still accepted.

**Figure 4.**
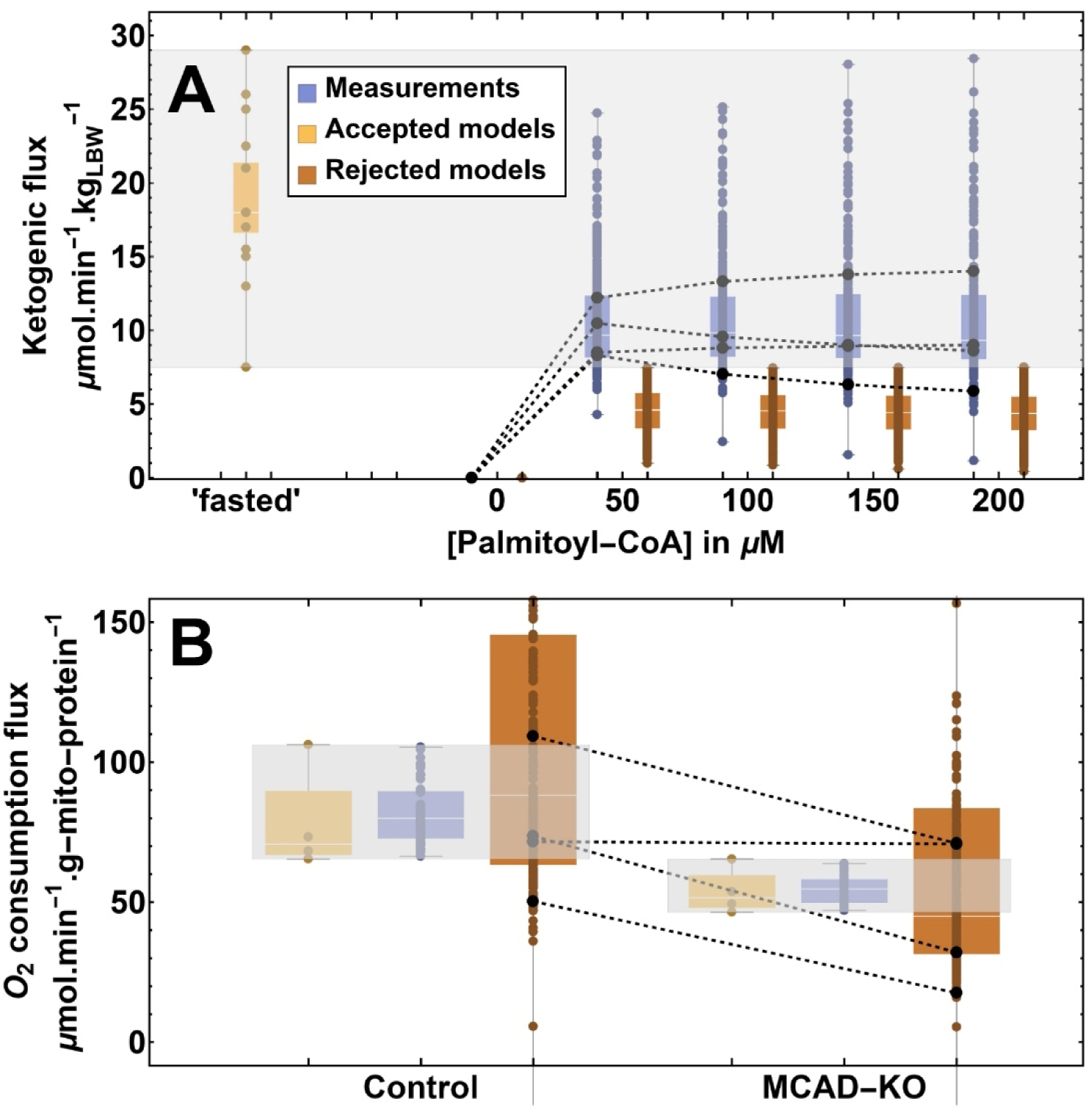
Model selection. **A.** First, the models were evaluated for their ability to predict the outcome of a whole-body ketogenic flux experiment [40]. If the model predicted a ketogenic flux within the range of measurements (grey) at any cytosolic palmitoyl-CoA concentration without ever passing above it, then the model was accepted and subjected to the next test. Four representative accepted models are explicitly shown as data points joined by dashed lines. In total, 585 models were generated of which 274 passed this first test. **B.** The 274 models that passed the first selection were tested for their ability to predict the O_2_ consumption flux by control and MCAD-knockout HepG2 cells at 25 µM palmitoyl-CoA [7]. If both fluxes were within the range of measured flux (grey boxes), then the model was accepted. The outcomes of four representative rejected models and one accepted model are shown in black and connected with dashed lines. In total 50 generated models passed this test, as well as the default model, giving 51 validated models.

A second experiment was then simulated: mitochondrial O_2_ consumption in the presence of palmitoyl-CoA. The O_2_ consumption flux of HepG2 cells, both wild-type and MCAD-knockout, was measured [7]. In the assay, a saturating concentration of palmitoyl-CoA was the only oxidative substrate provided, hence the O_2_ consumed during the full oxidation of palmitoyl-CoA could be calculated based on the known stoichiometry by which NADH and FADH_2_ are produced and used during respiration to produce ATP [44]. Furthermore, proteomics data were used to convert the default model, which is based on human hepatocytes, to a model that represents HepG2s, which were used in the experiments [45]. All conversions were done as in [7]. The simulation results from the 274 models that passed the first round of selection were then tested against the abovementioned experiments (Fig. 4B). Models were accepted if the predicted fluxes were within the range of measured values for both wild-types and knockouts.

The parameter values of the accepted ensemble are presented in Table S1 and the parameter values of the rejected ensemble in Table S2.

To assess which parameters had a strong effect on whether or not a model met the selection criteria, the parameter values in the accepted and rejected models were compared. Rejected models had a V_max,VLCAD_ lower than the range of all accepted models more than 65% of the time (Table S3).

No other parameter showed such a large difference between the rejected and accepted models even 10% of the time. For example, for V_max,MCAD_, 9.2% of rejected models had values higher than the range of values in the accepted models, while for V_max,SCAD_, 11% of the models had values lower than the range of accepted models. Both the lowest retrieved literature value for V_max,VLCAD_ (0.003 μmol.min^−1^.mg-mitochondrial-protein^−1^) [46] and the second lowest (0.0345 μmol.min^−1^.mg-mitochondrial-protein^−1^) [47] were below the range of values in the accepted models (Table S3). Only the highest retrieved literature V_max,VLCAD_ was within the range of the values observed in the accepted models. This draws the validity of the other two literature values in question (Fig. S15A; Table S3).

To evaluate this further, in each of the 51 accepted models the V_max,VLCAD_ was replaced by the respective literature values and subjected to the selection tests again (Fig. S15B-D). Where the lowest of the three V_max,VLCAD_ literature values (0.003 μmol.min^−1^.mg-mitochondrial-protein^−1^ [46]) was used, all previously accepted models were rejected. In contrast, the two highest values (0.0345 μmol.min^−1^.mg-mitochondrial-protein^−1^ [47] and 0.076 μmol.min^−1^.mg-mitochondrial-protein^−1^ [48]) led to 11 models each being reaccepted. The behaviour of the latter two adjusted ensembles was very similar (Fig. S15E-F), suggesting that even higher V_max,VLCAD_ values were generally present in accepted models.

### Ensemble simulations of ACAD deficiencies

To test the ensemble’s ability to describe the biochemical effects of mitochondrial fatty acid oxidation defects (mFAODs), we applied it to the case of MCAD deficiency (MCADD). Previously, the single-parameter default model was used to simulate the NADH production flux and mitochondrial acyl-CoA profile, including free CoA (CoASH) concentration, of an MCADD model *versus* a control model [7]. Some of this predicted behaviour, specifically, the increase in C8-acyl-CoA and the C8/C10 ratio that is used in clinical diagnosis, combined with a decrease in long- and short-chain acyl-CoAs, and CoASH, was experimentally validated in HepG2 cells [6]. The ensemble also recapitulated this behaviour, yielding a significant increase in C8-acyl-CoA and significant decreases in C4-CoA and CoASH (Fig. 5A), and a visible increase in the C8/C10 ratio in the MCADD relative to the control model (Fig. 5B).

**Figure 5.**
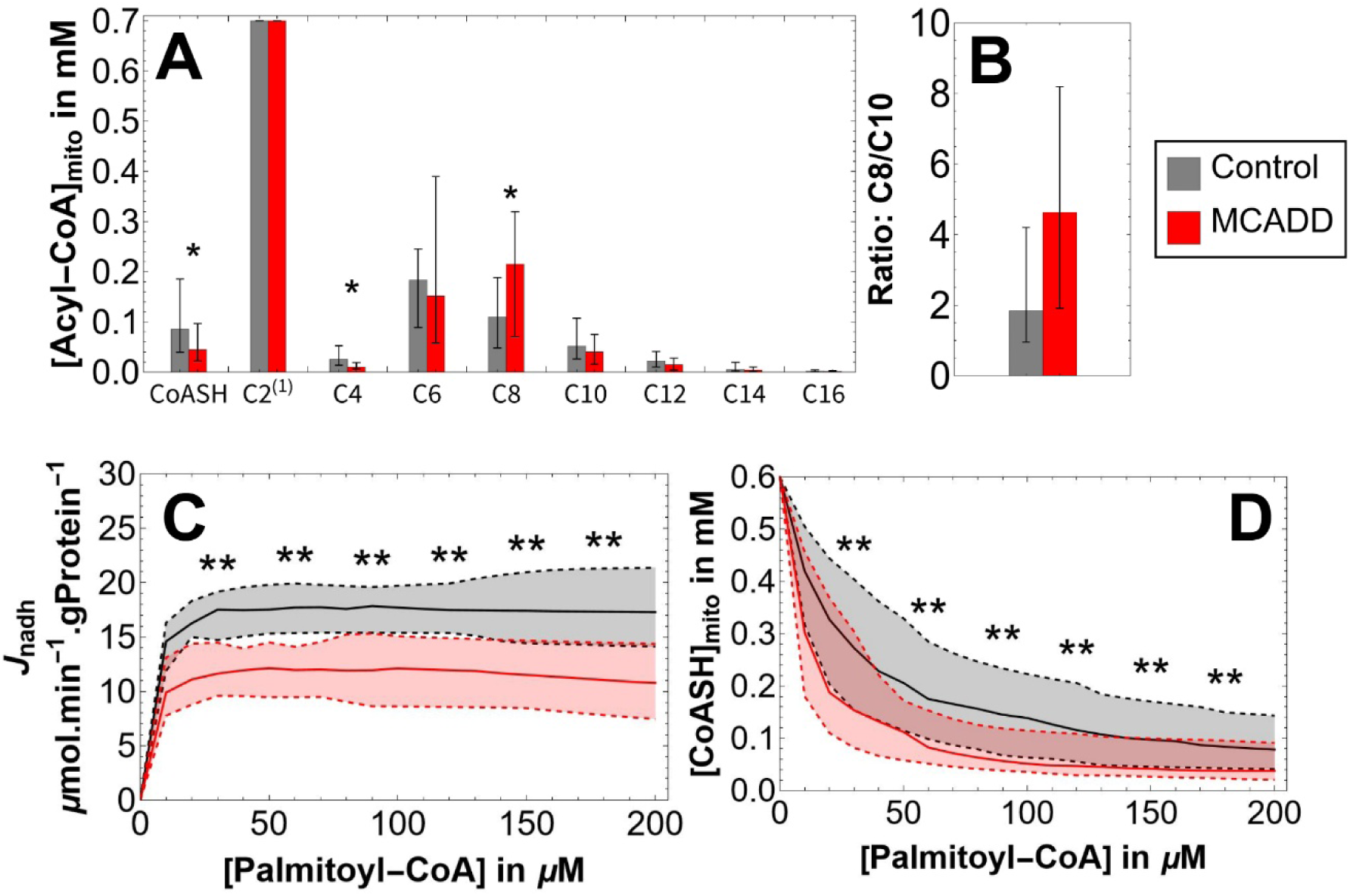
Ensemble prediction of the effect of MCADD. The shaded areas between the dashed lines indicate the interquartile range of the outcomes from the ensemble of simulations. The solid line represents the median. The ensemble without deficiency (Control) is compared to a model with zero MCAD activity (MCADD). Results are shown for the concentrations of the various mitochondrial acyl-CoA chain-lengths, including free CoA (CoASH) (**A**) and the C8/C10 ratio (**B**) at a cytosolic palmitoyl-CoA concentration of 150 μM, as well as the NADH production flux (J_nadh_, **C**), and the free mitochondrial CoA concentration (CoASH, **D**) at various cytosolic palmitoyl-CoA concentrations. Asterisks indicate significance of the difference between the Control and the MCADD ensembles using the Mann-Whitney U test (*p p < 0.01** p < 0.01). ^(1)^ C2 (acetyl-CoA) was fixed.

Next, the concentration of the substrate (cytosolic palmitoyl-CoA) was varied. An increasing substrate concentration reflects the fasted state, in which fatty acids are released from the adipose- tissue stores and supplied to the liver. Since NADH is one of the products of the mFAO, the NADH production flux was plotted as a measure of mFAO flux (Fig. 5C). MCADD deficiency significantly decreased the NADH production flux at all palmitoyl-CoA concentrations above 20 μM.

In line with a previously hypothesised phenomenon in which inborn errors of metabolism like ACAD deficiencies lead to the depletion of CoASH [9,49–51], we also investigated the CoASH concentration as a function of palmitoyl-CoA (Fig. 5D). MCADD significantly decreased the concentration of CoASH relative to the control model, in agreement with previous predictions from the single parameter model [7]. Indeed, later experimental work also indicated that CoASH was decreased in MCAD-knockout relative to control HepG2 cells [6]. This shows that the ensemble model replicates previous predictions about the dynamics of at least one mFAOD. Additionally, the application of the ensemble allows us to associate a statistical significance to the differences between control and mFAOD models, removing the arbitrariness and bias associated with parameter selection.

### Comparison of different FAODs

The ensemble was applied to a set of known mFAODs that could be simulated by the model. Table 1 shows a list of known mFAODs with some biochemical and physiological indicators, as well as an indication of the severity and association with hepatopathy and hypoglycaemia of each (cf. [52]). It was not the objective to present an exhaustive account of the literature associated with these mFAODs. Rather, any report of a clinically symptomatic individual with a given mFAOD, along with the reported residual enzyme activity, was used to define the approximate range of activity of each enzyme for which a given mFAOD might lead to a large enough disruption in the mFAO to cause hepatopathies or hypoglycaemia.

**Table 1.**
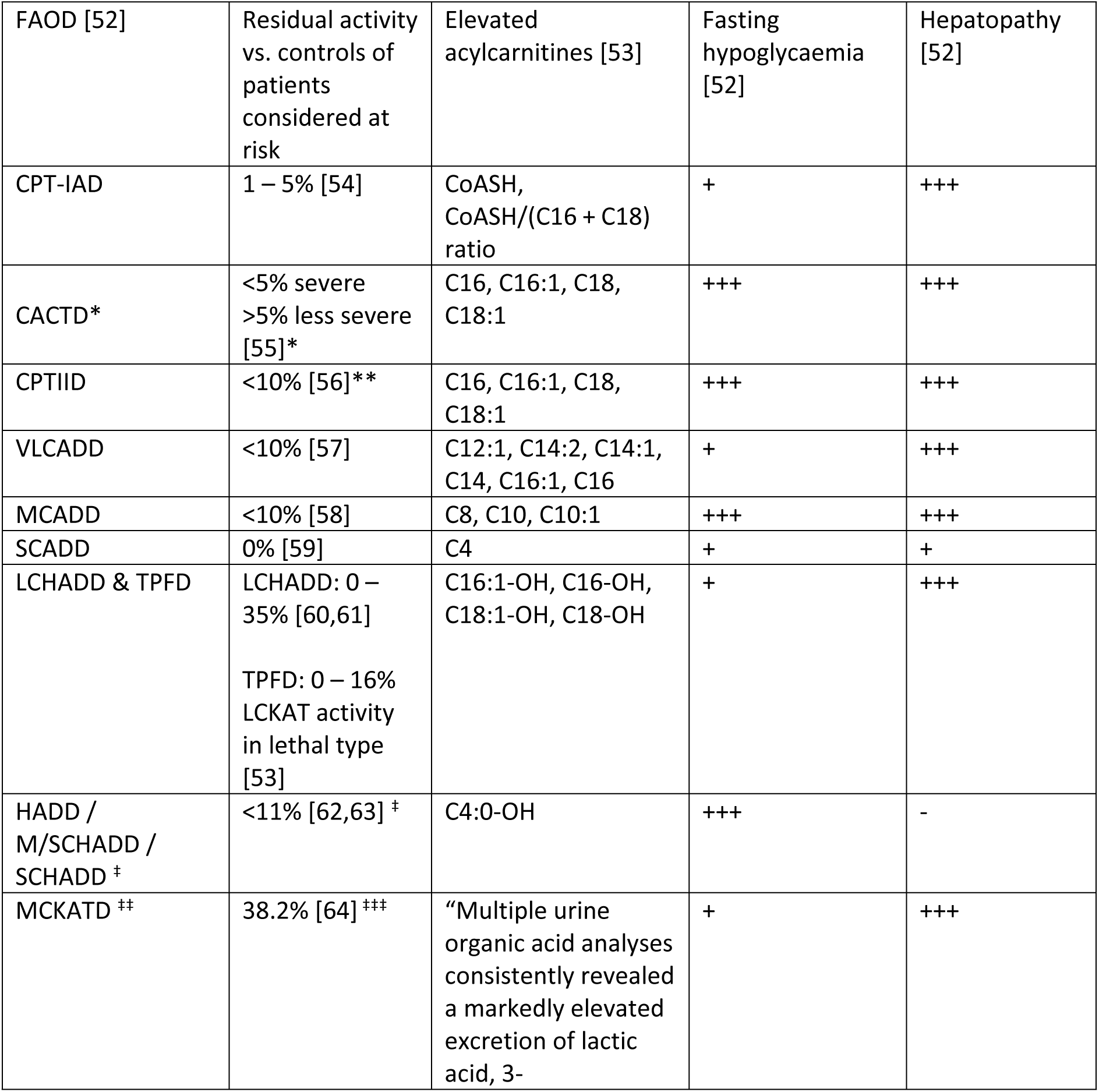
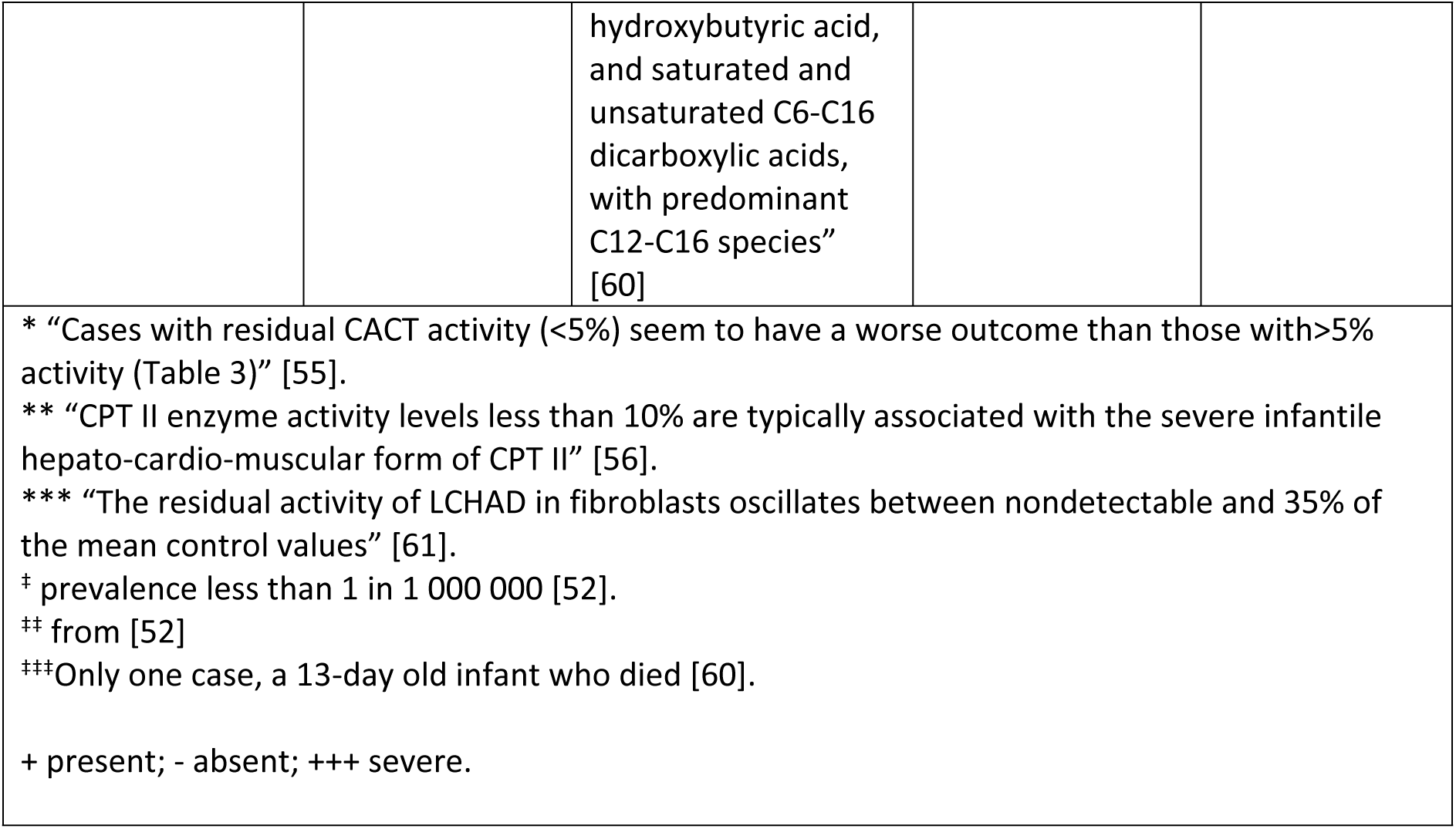
Biochemical and clinical characteristics of mitochondrial fatty-acid oxidation defects.

The V_max_ of each of these enzymes was incrementally decreased down to 0%. For all simulations, cytosolic palmitoyl-CoA was set to 150 μM to represent a stressed state at which one would be very reliant of the mFAO for energy homeostasis [7,65]. NADH production flux and mitochondrial CoASH concentration are shown as key readouts in these simulations (Fig. 6). Fig. 6A-J shows the effect of titrating down the activity of enzymes responsible for long-chain (LC-)mFAO. While the enzymes sometimes have a broad substrate-specificity *in vitro* [66], *in vivo* they are anchored to the mitochondrial membranes, which predisposes them even more to preferentially oxidizing the hydrophobic, membrane-bound long-chain substrates [67]. Likewise, Fig. 6K-R shows the effect of decreasing the activity of medium- and short-chain-specific (S/MC)-mFAO enzymes which are soluble, located in the mitochondrial matrix, and tend to interact more with the more hydrophilic, shorter intermediates that diffuse away from the mitochondrial membrane into the bulk solvent of the matrix [67].

**Figure 6.**
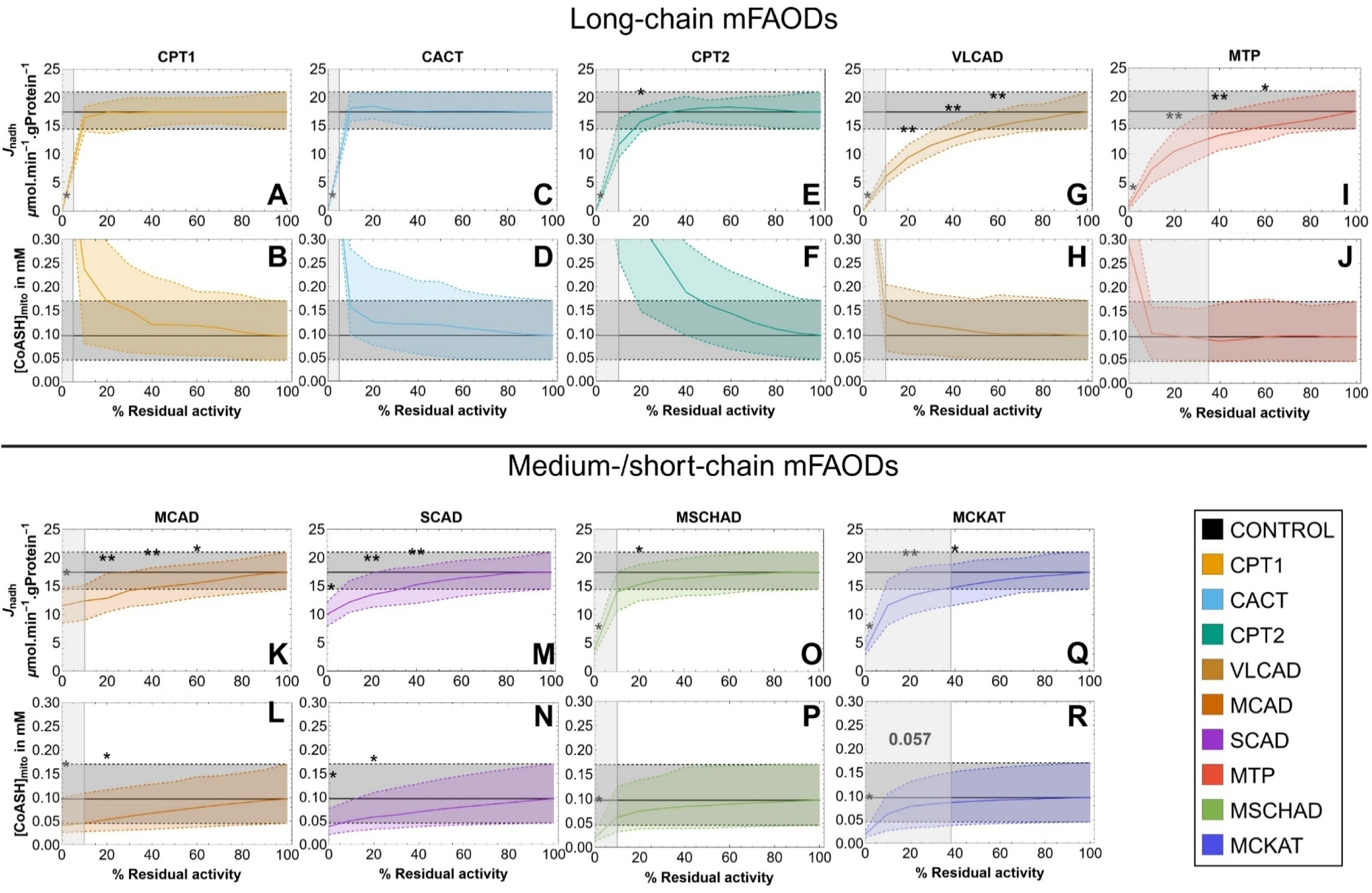
Long-chain mFAODs and medium-/short-chain mFAODs have different metabolic implications. NADH production flux (J_nadh_, in units of μmol.min^−1^.mg-mitochondrial-protein^−1^) and free mitochondrial CoA levels ([CoASH]_mito_) are compared for long-chain (LC)-mFAODs *versus* medium-/short-chain (S/MC-)mFAODs as a function of residual enzyme activity at 150 μM of cytosolic palmitoyl-CoA. The shaded vertical grey area on each figure represents a range of residual enzyme activities at which a person might be at risk, or at which symptoms have been reported (Table 1). The shaded areas between the dashed lines indicate the interquartile range of the outcomes from the ensemble of simulations. The solid line represents the median. The horizontal shaded area pertains to the ensemble without any enzyme deficiency (control). Results are shown for deficiencies of carnitine palmitoyltransferase 1 (CPT1, **A** & **B**), carnitine-acylcarnitine translocase (CACT, **C** & **D**), carnitine palmitoyltransferase 2 (CPT1, **E** & **F**), very long-chain acyl-CoA dehydrogenase (VLCAD, **G** & **H**), mitochondrial trifunctional protein (MTP, **I** & **J**), medium-chain acyl-Coa dehydrogenase (MCAD, **K** & **L**), short-chain acyl-CoA dehydrogenase (SCAD, **M** & **N**), medium- /short-chain hydroxyacyl-CoA dehydrogenase (MSCHAD, **O** & **P**), and medium-chain ketoacyl-CoA thiolase (MCKAT, **Q** & **R**). Asterisks indicate significance of the difference between the control and the deficiency according to the Mann-Whitney U test (* p < 0.05, ** p < 0.01).

LC-mFAODs have a very acute, statistically significant effect on the pathway flux, especially at very low (<10%) levels of residual activity (Fig. 6A, C, E, G, I). This differs according to the specific LC-mFAOD, for instance with decreases in VLCAD and MTP residual activity having a more gradual effect on pathway flux (Fig. 6E & G), though all LC-mFAODs tend to zero flux at zero residual activity.

Importantly, the range of residual enzyme activities associated with disease roughly corresponds to the range in which the flux drops significantly below control flux. Mitochondrial CoASH concentrations are not depleted; rather, at the lowest residual activities, they go up, because the lack of substrates entering the pathway implies that very little CoA is acylated (Fig. 6B, D, F, H, J).

S/MC-mFAODs have a qualitatively different effect: both flux and CoASH decrease with decreasing residual activity (Fig. 6K, M, O, Q). In contrast to LC-mFAODs the flux does not decrease to zero. CoASH concentration decreases as a function of residual activity for all S/MC-mFAO enzymes, with significant differences detected at the lowest residual activities for all S/MC-mFAODs (Fig. 6L, N, P, R). There is no unequivocal correlation between the residual activity below which clinical symptoms occur on the one hand and the decrease of flux or CoASH on the other hand (Table 1). This indicates that caution should be exercised when translating these results to clinical applications.

## Discussion

We set out to construct a model with which we could explore the dynamics underlying various mitochondrial fatty acid oxidation defects (mFAODs). In doing so, we discovered ample variation in the values reported for kinetic parameters. Such parameter uncertainty is seldom explicitly considered in kinetic models of metabolism. Rather, the best or – more concerningly – most convenient parameter is often chosen, and any considered alternatives discarded [68]. Often the justification for this might be that small differences in parameter values likely do not have a large effect on model behaviour. Contrast this, however, with Saa and Nielsen’s [24] observation that “differences in reported data can be as great as three orders of magnitude” – hardly a small deviation. Indeed, in our parameter search, equally plausible assays – evaluated to the best of our biochemical knowledge – often yielded variability exceeding an order of magnitude (Fig. 2).

Moreover, experimental variability is especially large for human-derived kinetics, much more so than in more controlled studies of mice or rats [17]. Human samples cannot easily be subjected to rigorously standardised assay conditions or interventions. Most primary human material is obtained from discarded or partial transplants, or from excised tissue [16]. The scarcity of primary human material means that kinetic information from individuals of different ages, sexes, and ethnic and medical backgrounds are often grouped [17]. Moreover, methodological differences like tissue procurement and processing further contribute to variability [69]. Uncertainty can therefore be biological or technical in origin – or both [70].

Against this background, we decided to build a computational kinetic model of the mitochondrial β-oxidation in human liver which incorporates uncertainty about kinetic parameters. We are not aware of any other models of the fatty acid β-oxidation that explicitly incorporate kinetic uncertainty. The ensemble allowed us to reduce the arbitrary bias of fixed-parameter models in which plausible alternatives are discarded [68]. Variation was constrained based on underlying patterns among chain length-specific parameters, which has not been done in previous ensemble kinetic modelling approaches [18,33,68,70]. A model selection step ensured that the ensemble contains only models that make predictions within the range of measured data. Given the large number of model parameters and their sometimes excessively large variation, this approach allowed optimal use of all data, while still acknowledging substantial parameter uncertainty and incorporating this into our predictions. 534 models were rejected before 50 were accepted in addition to the default, yielding an ensemble of 51 models.

A high V_max,VLCAD_ value was strikingly important for a valid model: the rejected models had lower V_max,VLCAD_ values than the accepted ensemble in 66% of the cases. Testing all model parameters that were free to vary during ensemble selection, no other parameter showed similar differences between the rejected and accepted ensembles in a particular direction more than 15% of the time. Table S3 shows that the deviation is much less pronounced for V_max,SCAD_ and V_max,MCAD_, for example. Moreover, substituting the lowest literature-derived V_max,VLCAD_ value back into the accepted ensemble caused all previously valid models to miss the selection criteria. This measurement was performed on human foetal tissue fixed with 4% paraformaldehyde, embedded in paraffin [46], and subsequently shipped between laboratories (*personal communication*). This handling will have destroyed much of the enzyme activity in the sample, rendering it inappropriate as a measure of the absolute activity [69]. The fact that, out of 369 model parameters, only the lowest measurement of V_max,VLCAD_ correlated with poor performance against the selection criteria, suggests either that the distributions of measurements for other parameters are within the correct range, or lack control over metabolic flux.

After construction and validation, we applied the ensemble to the case of MCAD deficiency (MCADD). We have previously used the default model to explore the biochemical dynamics underlying the occurrence of hypoketotic hypoglycaemia in MCADD, suggesting that the combination of CoASH depletion and a moderate flux impairment might underlie these metabolic decompensations [7]. More recent work, using MCAD-knockout HepG2 cells, has confirmed that medium-chain acyl-CoAs accumulate under conditions of energetic stress, leading to a decline in CoASH [6]. The ensemble recapitulated those results, lending credence to its utility as a tool for investigating mFAODs (Fig. 5).

The ensemble was then applied to a set of known mFAODs (Table 1). We subdivided these mFAODs into long-chain mFAODs (LC-mFAODs) and short-/medium-chain mFAODs (S/MC-mFAODs). LC-mFAODs had a very acute effect on pathway flux, and it was striking to observe that the ranges of residual enzyme activity values typically associated with clinical symptoms corresponded to the ranges in which the flux differed significantly between the mFAOD and control ensembles (Fig. 6A-J). VLCAD deviated most from this trend, with significantly lower flux at higher residual VLCAD activities than the associated literature range. However, pathological cases of VLCADD with residual activity as high as 22% are known, despite being regarded as less consistently diagnostic [57].

This association of LC-mFAODs with pathway flux echoes the observation that the extent of LC-FAO flux impairment in VLCAD deficient fibroblasts predicted disease severity in patients [71]. The strong association of LC-mFAODs with flux might also explain the recent success of triheptanoin administration to patients with LC-mFAODs [72]. Triheptanoin, a triglyceride containing three heptanoates yields two acetyl-CoAs and one propionyl-CoA as final products. Propionyl-CoA can be converted to succinyl-CoA, which can enter the TCA cycle and augment ATP production, both of which are necessary for gluconeogenesis. It is known that TCA cycle intermediates are depleted in LC-mFAOD patients, and that ATP production is decreased [73]. Both can be seen as effects of deficient mFAO pathway flux, as acetyl-CoA, the terminal product of the mFAO, is produced more slowly when pathway flux is reduced. Moreover, as gluconeogenesis anaplerotically consumes TCA cycle intermediates in the liver, the TCA cycle’s intermediates can become depleted if sufficient mFAO flux is not sustained [74]. Triheptanoin provides a way of restoring some TCA cycle flux without having to pass via all the LC-mFAO reactions.

S/MC-mFAODs, on the other hand, displayed a combination of flux-impairment – though not as severe as LC-mFAODs – and a reduction of CoASH (Fig. 6K-R). This echoes our own recent work on MCADD [6,7]. CoASH depletion, a fundamental component of the so-called CoA sequestration, toxicity and redistribution (CASTOR) effect [49,51,75] has recently also been related to propionic acidaemia [76,77]. CoASH depletion can have pleiotropic effects, as it is not only an important substrate for import of mFAO substrates into the mitochondrion via CPT2 [78], but also a substrate for a number of other pathways, including the TCA cycle [79,80] and branched-chain amino acid oxidation [81].

It is striking that, in the model, all S/MC-mFAODs cause a reduction of CoASH, while severe LC-mFAODs rather lead to increases in CoASH. S/MC-mFAODs affect enzymes in the middle or at the end of the ß-oxidation spiral. This means that there are many more metabolites upstream of them, which all sequester CoA. The decrease in CoASH is also particularly strong when including the effect of metabolite partitioning– a model a model attribute which imitates the effect of the decreased water solubility of long-chain intermediates [67] (compare Fig. 6 *versus* Fig. S16). Long-chain pathway intermediates tend not to leave the hydrophobic membrane environment but remain anchored into the membrane with their hydrophobic tails [65,82,83]. This increases their effective concentrations in the hydrophobic environment of the membrane-bound LC-mFAO enzymes [67]. These enzymes are then much closer to saturation at lower acyl-CoA concentrations, which generates a high influx of short- and medium-chain pathway intermediates that sequester CoA. The effective concentrations of short- and medium pathway intermediates are closer to their real concentrations, so they accumulate more before saturating their enzymes, leading to more CoA sequestration. These accumulations are exacerbated by S/MC-mFAODs, which reduce the capacity of the downstream enzymes to oxidise these accumulating intermediates. LC-mFAODs, however, are situated upstream of these accumulations, thereby alleviating CoA sequestration.

In this work, we have presented an ensemble kinetic model of human hepatocytic mFAO. The 51 models constituting the ensemble were selected for their ability to predict certain known biochemical readouts. Simulation of the effect of MCADD echoes what was recently seen experimentally i.e., that medium-chain acyl-CoAs accumulate, decreasing the size of the CoASH pool, which is required in many mitochondrial pathways. Finally, we explored the biochemical effects of a set of mFAODs, subdivided into LC-mFAODs and S/MC-mFAODs, finding that LC-mFAODs are flux-impairment diseases associated with decreases in residual enzyme activities. S/MC-mFAODs, on the other hand, are more complex, with simultaneous flux and CoASH sequestation effects. This work can help aid researchers and clinicians to understand the relationship between different mFAODs better, perhaps contributing to better risk stratification and treatment development.

## Methods

### Software

All computation was carried out using Wolfram Mathematica ® version 13.1. Wherever functions are mentioned, it pertains to built-in function in Mathematica ®.

### Default model

odendaal4, a published kinetic model of the mitochondrial fatty acid oxidation as in occurs in human liver, was used as the starting point for this study [6]. It was based on Michaelis-Menten kinetics and experimentally determined *in vitro* enzyme kinetics. One adjustment was made, for comparability across the other parameter sets: instead of modelling acetyl-CoA as a variable which is exported by mass action kinetics, it was set to a fixed value of 700 μM [84]. This model is referred to as the “default model” throughout this work.

### Literature search

Kinetic parameters were retrieved from the literature or calculated or inferred based on literature-derived information. The enzyme name and Enzyme Commission Number (EC number) were used as search terms on the Braunschweig Enzyme Database (BRENDA; [85]) and SABIO-RK [86]. For CACT, which does not have an EC number, the UniProtKB primary accession number (O43772) was used.

The enzyme name and the words “liver” and “kinetics” were also searched on Google Scholar and MEDLINE. The first 200 entries were considered and judged, based on their titles, for their applicability. Abstracts of articles containing the words “characterisation”, “kinetics”, “human”, or the enzyme name in the title were read. Based on the abstract a decision was made whether or not to read the whole article.

Enzyme kinetics was retrieved from these articles, including pH-, temperature-, and buffer composition-dependence, tissue-specificity, and membrane-embedding. All relevant information was noted in a report per enzyme or boundary condition.

Laboratory measurements of human liver kinetic parameters could not always be found. Where satisfactory human parameters were not available, values from other mammals were also considered. Where satisfactory liver parameters could not be found, other tissues were also considered. Where no appropriate values could be found, they were assumed from similar enzymes, inferred from related parameters, or calculated by the Haldane relation [87]. These are available on FAIRDOM Hub [88] (DOI: 10.15490/fairdomhub.1.assay.2137.2).

### Weighting parameters

A weight between 0 and 1 was assigned to each parameter value to express the confidence that it reflects the *in vivo* environment. For consistency, one of four weight categories was assigned in each case: 0.1 (very implausible), 0.5 (uncertain), 0.9 (plausible) and 1.0 (very plausible). A parameter was usually penalised by downgrading it one category for each assay condition that deviated from physiological environment. The criteria are enzyme-specific (DOI: 10.15490/fairdomhub.1.assay.2137.2). Temperature, pH, organism, and tissue of origin were always considered.

### Constructing the distributions

For chain length-dependent values, like K_m_ and sf values, multinormal distributions were constructed. Equations were fitted to the parameters as a function of acyl chain length. The equations were mostly cubic polynomials (Fig. 3A & B), but also included parabolic, exponential, and linear functions. The fitted function and its covariance matrix were then inserted as variables into Mathematica®’s *MultinormalDistribution* function.

For chain length-independent parameters, which include V_max_ values, inhibition constants (K_i_), and chain length-independent K_m_ values, univariate distributions were constructed. Following the suggestion of Tsigkinopoulou *et al.* [18], log-normal distributions were made (using the *LogNormalDistribution* function), unless a lack of good data made it impossible to determine appropriate parameters for a log-normal distribution. Otherwise, normal or uniform distributions were assigned using the *NormalDistribution* function. Standard deviations in normal distributions were set to 25% of the mean. Uniform distributions were assigned ranges from which values were randomly selected using the *RandomReal* function.

The distributions and their statistical parameters, covariances, as well as the values used to generate them are shown per kinetic parameter in Text S1. Where appropriate, the functions fitted to chain length-dependent parameters are also presented with their coefficients and plotted next to the literature data for comparison.

The visualisation of probability density function for the comparison of the rejected and accepted parameter space in Fig. S15A-E was done by feeding the data to Mathematica®’s built-in *PDF* function.

### Generating parameters

Following Achcar *et al.* [70], parameter values were sampled from the distributions to generate an ensemble of plausible models. In the interest of model tractability, equilibrium constants (K_eq_), boundary conditions, and the kinetics of ACOT were kept constant. Five parameters were calculated based on the ensemble of other parameters generated for a reaction: Kicpt1CarCYT, Kicpt2CarMAT, KmmckatAcylCoAMAT, KmmtpAcylCoAMAT, and Vrcact. The first two were equal to the mean K_m_ of the enzyme for all across chain lengths, the latter three were calculated according to the Haldane relation [36]. The first two K_i_ values are chain length-independent parameters related to chain-length-dependent K_m_ values. They were assumed as the mean of the K_m_ values, as was the case in the fixed-parameter model [6,7]. The latter three parameters are dependent on other parameters via the Haldane relation [36]. The solution for this calculation for KmmckatAcylCoAMAT is shown in Fig. 3F.

Other parameters were sampled from their distributions. For uniform distributions, a range was passed to Mathematica®’s *RandomVariate* function. For normal, log-normal, and multinormal distributions the constructed distribution was passed to *RandomVariate* to sample parameter values, either per chain length or independent, as the case may be. Bounds were placed on the possible parameter value by rejecting parameters that do not fall with in the permissible range. These ranges were chosen to best reflect the variation in the data and are presented in Text S1. Ten-fold higher and ten-fold lower than the chosen parameter from the fixed-parameter model was selected as the bound in >95% of the cases [6,7]. A representative set of outcomes are presented next to the literature data for all parameters in in Text S1.

### Thermodynamic consistency

Thermodynamic consistency was ensured by sampling parameters from rate equations that contained a K_eq_ instead of the ratio between forward and reverse V_max_ values [70]. The transporter, CACT, explicitly contains the forward and reverse V_max_. Thermodynamic consistency was ensured by calculating the reverse V_max_ using the Haldane equation [36].

### Calculating steady states

Time evolution simulations were carried out with all variable concentrations set to starting values of zero until apparent steady states were reached. This was carried out using Mathematica®’s *NDSolve* function. The default method selection function (*method → “Automatic”*, in Mathematica® code) was used in all cases except for the results in Fig. S15, in which stiffness necessitated the explicit use of the backward differentiation formula (*method → “BDF”*). The apparent steady states calculated in this way were then used as initial guesses in the *FindRoot* function, in which the real steady state was calculated by setting all ODEs == 0, always using Mathematica®’s default method selection function (*method → “Automatic”*).

### Model simulation

All simulations were carried out using the ensemble parameter values in Table S1 (accepted ensemble parameters) or Table S2 (rejected ensemble parameters). All deviations from these values are explicitly indicated in figure legends or Model Selection in the Methods section.

### Model selection

Each model was tested for its ability to simulate two sets of data: whole body ketogenic flux as reported by Fletcher *et al.* [40] and a HepG2 oxygen consumption flux (wild type and MCAD-knockout) measured by us and reported in a previous publication [36]. Model adjustments were carried out exactly as in a previous publication, in which the same data were used for model validation [36].

For whole-body ketogenic flux, each generated model had to predict a flux that surpassed the minimum of the range at one of the investigated palmitoyl-CoA concentrations without every surpassing the maximum. For the oxygen consumption, both the wild-type and MCAD-KO models had to predict a flux within the range of measured values. Only of all of these criteria were met, was a model accepted as part of the ensemble.

### Statistics

Ensemble predictions were statistically compared in using the Mann-Whitney U test, as prediction ranges were not normally distributed. This was done using Mathematica®’s built-in function *MannWhitneyTest*.

## Data availability statement

Data is stored and made available on FAIRDOM Hub: a repository and collaboration environment for sharing systems biology research. They form a part of a larger investigation (DOI: 10.15490/fairdomhub.1.investigation.587.2). The relevant subfolders via which the supporting documentation is available are found at: kinetics minireviews used to evaluate and weight parameters (DOI: 10.15490/fairdomhub.1.assay.2137.2) and the notebooks and data files used to generate the ensemble, run simulations, and plot the results (DOI: 10.15490/fairdomhub.1.study.1460.1).

## Acknowledgements

The authors would like to thank José M. Horcas Nieto, Ligia A. Kiyuna, Emmalie A. Jager, Karen van Eunen, and Marcel A. Vieira Lara for reading, correcting, and supplementing the mini-reviews from which enzyme kinetics were extracted. Dirk-Jan Reijngoud, Fentaw Y. Abegaz, Oliver Ebenhöh, and and Marcel A. Vieira Lara also contributed substantially through stimulating discussions. Figures 1 and 2A were created with BioRender.com. This work was supported by the European Union’s Horizon 2020 research and innovation programme under the Marie Skłodowska-Curie Actions Grant Agreement PoLiMeR, No 812616 and by the University Medical Centre Groningen.

## Supplementary Materials

Figure S1. CPT1 parameters.

Figure S2. CACT parameters.

Figure S3. CPT2 parameters.

Figure S4. CrAT parameters.

Figure S5. VLCAD parameters.

Figure S6. MCAD parameters.

Figure S7. SCAD parameters.

Figure S8. CROT parameters.

Figure S9. MSCHAD parameters.

Figure S10. MCKAT parameters.

Figure S11. MTP parameters.

Figure S12. ACOTci parameters.

Figure S13. ACOTcs parameters.

Figure S14. Boundary conditions, conserved moiety sums, compartment volumes, and the percentage of CPT located at membrane contact sites.

Figure S15. Feasibility of measured V_max,VLCAD_ values.

Figure S16. Long-chain mFAODs and medium-/short-chain mFAODs without metabolite partitioning.

Text S1. Parameter values, weights, and distributions.

Table S1. Ensemble parameters.

Table S2. Rejected ensemble parameters.

Table S3. Parameter sensitivity.

